# The efficacy of antiseptic agents against staphylococcal biofilm can be quantitatively assessed with the use of processed microscopic images

**DOI:** 10.1101/2021.11.30.470646

**Authors:** Grzegorz Krasowski, Paweł Migdał, Marta Woroszyło, Karol Fijałkowski, Grzegorz Chodaczek, Joanna Czajkowska, Bartłomiej Dudek, Joanna Nowicka, Monika Oleksy-Wawrzyniak, Bartłomiej Kwiek, Justyna Paleczny, Malwina Brożyna, Adam Junka

## Abstract

The staphylococcal biofilms are major causative factors of non-healing wound infections. Their treatment algorithms recommend the use of locally applied antiseptic agents to counteract the spread of infection. The efficacy of antiseptics against biofilm is assessed *in vitro* by a set of standard quantitative and semi-quantitative methods. The development of software for image processing allowed to obtain quantitative data also from microscopic images of biofilm dyed with propidium iodine and SYTO-9 reagents, differentiating dead cells from the live ones. In this work, the method of assessment of the impact of antiseptic agents on staphylococcal biofilm *in vitro*, based on biofilm’s processed images, was proposed and scrutinized with regard to clinically-relevant antiseptics. Taking into account the fact that i*n vitro* results of the efficacy of antiseptic agents against staphylococcal biofilm are frequently applied to back up their use in hospitals and ambulatory units, our work should be considered an important tool providing reliable, quantitative data with this regard.

## 1. Introduction

The biofilms are aggregated communities of microbes, embedded within an extracellular matrix, consisting of various molecules (carbohydrate polymers, exocellular DNA, proteins). The biofilm is considered highly tolerant to antiseptics, antibiotics and immune system, not only due to the protective functions of matrix, but also due to the high concentration of cells within a relatively small surface and the differentiation of the cells’ metabolism/growth with regard to their spatial location **[1]**. The biofilms are major causative factors of numerous chronic diseases, including non-healing wounds. Treated improperly, they increase the risk of limb amputation or even the patient’s death **[2]**. The staphylococci (*Staphylococcus aureus*, mainly) are bacteria of particularly high ability to cause biofilm-based infections of non-healing wounds **[3]**. The present algorithms of treatment of these disease entities recommend (together with the debridement and application of dressings), the use of antiseptics, which are locally-applied antimicrobial liquids **[4,5,6]**.

The first stage of assessment of antiseptics’ efficacy against biofilm-forming pathogens involves *in vitro* analyses. Their performance (together with toxicity studies) are necessary, before any antiseptic can be used in the clinical conditions. For these *in vitro* analyses, quantitative culturing of staphylococcal cells and colorimetric (semi-quantitative) methods are mostly applied. The data obtained by these methods are frequently supported by microscopic images (of qualitative character) from confocal/epifluorescence microscopy, in which viable and compromised biofilm-forming cells are visualized with, among others, “Live/Dead” dyes **[7,8,9]**.

The emergence of computational software for image processing allowed to extract the quantitative data from such types of microscopic images **[10]**. The aforementioned Live/Dead dyeing, visualized with the use of microscopy and analyzed with image processing software is presently considered a reliable toolbox for determination of antimicrobial (and anti-biofilm) efficacy. Its level should correlate with an increase in the red fluorescence signal from the Propidium Iodide dye (PI, “Dead”), binding to non-altered cells, and/or drop in the green fluorescence signal from the SYTO-9 dye (“Live”), binding to the altered (damaged) cells. Such changes, recorded by means of image processing software, can be further extracted into numerical values and analyzed with the use of statistical methods **[11]**.

Nevertheless, the impact of phenomena taking place during biofilm culturing, introducing of antiseptic and dyeing of the biofilm samples, is frequently not taken under account in analysis of the image’s quantitative data on antibiofilm efficacy. It leads to discrepancies of outcomes obtained and use of biofilm images as a sort of ornamental results (supporting quantitative data from culturing or colorimetric methods). In turn, we hypothesized that by analysis of these phenomena and application of image processing software, the information on such crucial parameters of biofilms treated with antiseptic as change of their confluency, thickness, ratio of live to dead cells can be properly extracted and be perceived as data of at least equal value to data obtained by aforementioned classical methods.

Therefore, we decided to solve the above-mentioned issues and to increase the usability of confocal/epifluorescent microscopy in biofilm studies by provision of formula which allows to use the data from microscopic images and to calculate the antibiofilm effectiveness of the given antiseptic. Specifically, the three antiseptic products, broadly applied in the clinical setting, were used in this research - polyhexanide, povidone-iodine and hypochlorous solution, because of their different mechanism of action toward biofilms and biofilm-forming cells **[12,13]**.

Being fully aware of the fact that an excessive number of variables included in the analysis may lead to data overload, resulting in their misinterpretation, we decided to scrutinize only staphylococcal biofilm (and not biofilm of other potentially relevant species) and its tolerance/sensitivity to these three antiseptics. Therefore, the two specific goals of the present study was to analyze the phenomena occurring in the range of microscopic and molecular interactions during biofilm culturing, introduction of antiseptic and dyeing with L/D, and to use this knowledge to establish the way for a valid assessment (by means of microscopic and image processing methods) of the impact of antiseptics on biofilm *in vitro*.

## 2. Materials and Methods

### 2.1. Antiseptics applied

a. Prontosan wound irrigation solution® (B. Braun, Melsungen, Hessen, Germany), composed of purified water, 0.1% betaine surfactant, and 0.1% polyaminopropyl biguanide (polyhexanide), later referred to as PHMB;
b. Granudacyn® Wound Irrigation Solution (Molnlycke Health Care AB, Göteborg, Sweden), composed of water, sodium chloride, 0.005% sodium hypochlorite, and 0.005% hypochlorous acid, later referred to as NaOCl;
c. Braunol® (B. Braun, Melsungen, Hessen, Germany), composed 7.5% povidone-iodine with 10% available iodine, sodium dihydrogen phosphate dihydrate, sodium iodate, macrogol lauryl ether, sodium hydroxide, and purified water, later referred to as PVP-I.

### 2.2. Staphylococcal strains and biofilm culturing *in vitro*

Two reference strains from the American Type Culture Collection (ATCC), *Staphylococcus aureus* 6538 and 33591, and 8 clinical strains isolated from chronic wound infections were chosen. All staphylococcal strains are part of collection of the Department of Pharmaceutical Microbiology and Parasitology of Wroclaw Medical University, collected during research project approved by the Bioethical Committee of Wroclaw Medical University, protocol # 8/2016. The strains were transferred from Columbia Agar (Biomaxima, Lublin, Poland) to liquid Tryptic Soya Broth (Biomaxima, Lublin, Poland) and incubated for 24h/37°C. Next, they were diluted using densitometer (Densilameter II, Erba Lachema, Brno, the Czech Republic) to 0.5 McFarland and with serial dilution to ca. 10^5^ cfu/mL. The number of cfu was additionally checked by quantitative culturing on Columbia Agar. 2mL of staphylococcal suspensions of 10^5^cfu/ml were transferred to the wells of the 24-well plates (Wuxi Nest Biotechnology, Wuxi, China) and incubated for 24h/37°C. After that, the medium containing planktonic cells was gently removed, and the well was rinsed once with saline (Stanlab, Lublin, Poland). Such prepared biofilms were subjected to subsequent analyses. Above methodology applied in all cases with except:

– setting presented in **Figure 1A**, where biofilm was cultured for 12 hours

**Fig. 1.**
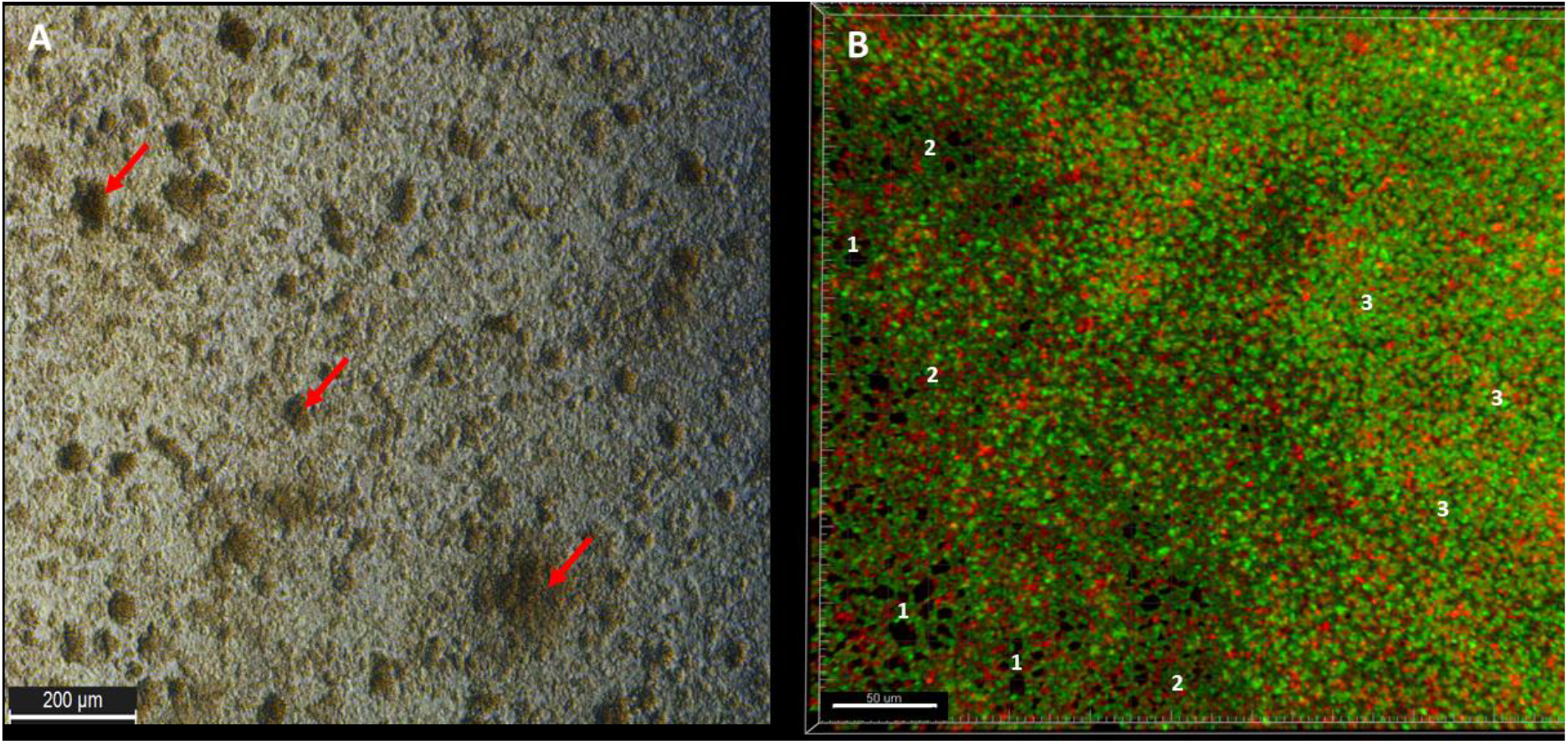
A – Young, 12h staphylococcal ATCC 6538 biofilm grown in a microplate (24-well) seen in an optical microscope (aerial perspective) without any prior preparative procedures; red arrows indicate areas of higher cellular density (mushroom-like structures); **B** – aerial perspective of L/D-dyed, 24h biofilm of the same strain. The “1s” show areas non-covered with cells (holes); the 2s” show areas of lower cellular density (of lower intensity of green color) than areas designated with “3s” (of higher density of green color). Red shapes are cells of decreased viability (with a compromised cell wall) or dead; while green shapes are viable cells (of non-compromised cell wall). Picture A, inverted microscope DMIL LED, magn. 10x; Picture B, confocal microscope SP8, magn. 40x.
– setting presented in **Figure 2** where biofilm was cultured for 24 hours but inside of microscopic glass (Medical Depot, Warsaw, Poland).

**Fig. 2.**
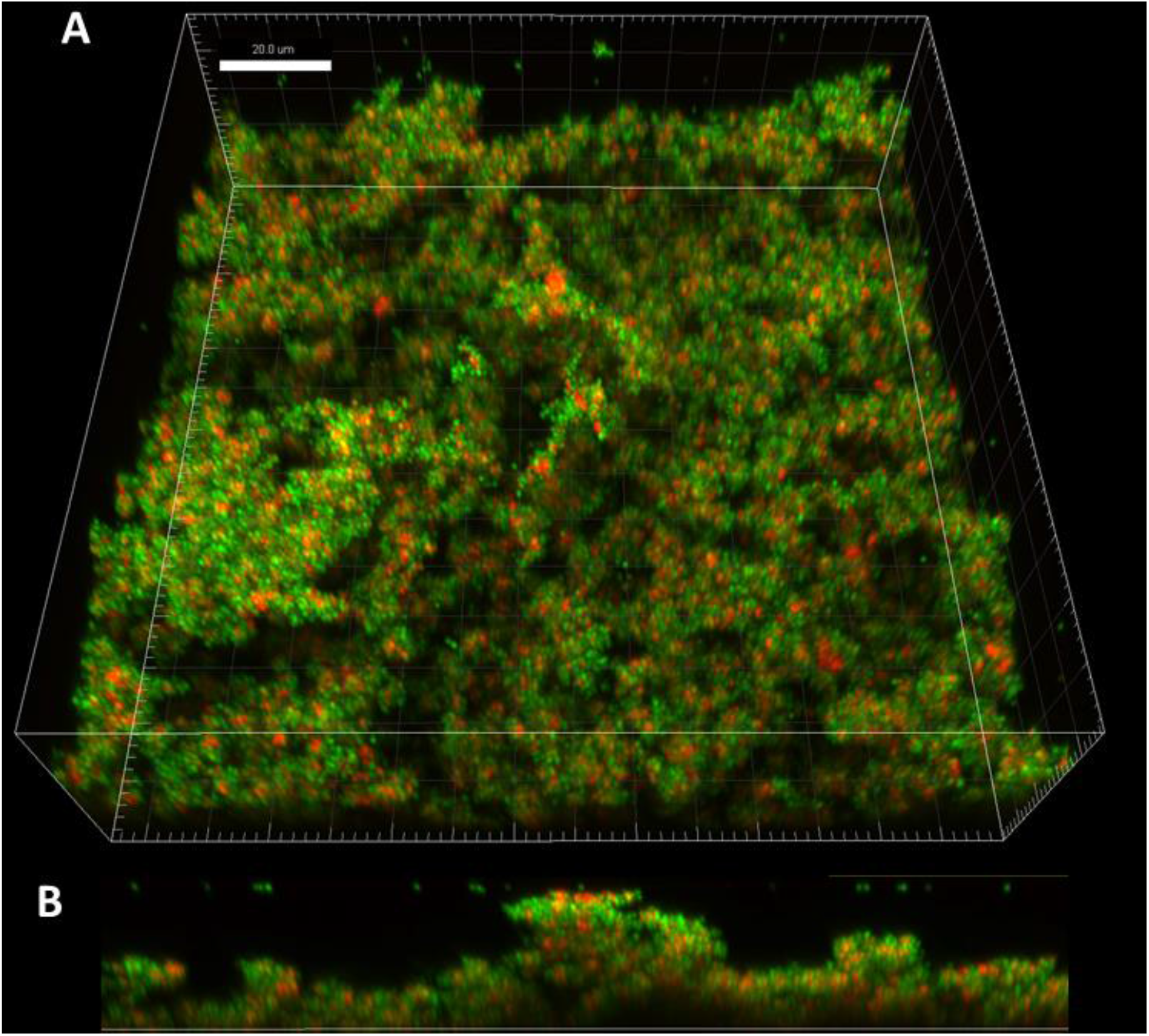
The Z axis image stack visualizing ATCC 6538 staphylococcal biofilm from the side aerial perspective (**A**). Part **B** of the figure shows the vertical cross-section through the three-dimensional biofilm structure, where the regions of various height and density are explicitly visible. Microscope Cell Observer, magnification 40x.
– setting presented in **Figure 6**, where medium was removed from 24-old biofilm and the images were taken using bright light vision in Etaluma Lumascope 620 fluorescent microscope (San Diego, CA, USA)

### 2.3. Live/Dead dyeing of staphylococcal biofilms and microscopic visualization

Staphylococcal biofilms, cultured as presented in Materials and Methods 2.2. were labeled with 500 µL of Live/Dead solution (Filmtracer™ Live/dead™ Biofilm Viability Kit, Thermo Fisher Scientific, Waltham, MA, USA) according to the protocol provided by the manufacturer. After 20 minutes of incubation without light, the staining solution was gently removed and samples were rinsed once with filter-sterilized water. Such prepared samples were subjected to subsequent analyses. The microscopic visualizations were performed using a wide-field LS620 fluorescent microscope (Etaluma, San Diego, CA, USA) and two confocal microscopes: a Cell Observer SD spinning disk system (Zeiss, Oberkochen, Germany) and an SP8 MP laser-scanning confocal microscope (Leica, Wetzlar, Germany). SYTO-9 showing live bacteria was excited at 488 nm wavelength using a LED (LS620) or a laser line (Cell Observer and SP8). The collected emission was within 502-561 nm (LS620), 502-538 nm (Cell Observer) or 500-530 nm (SP8) ranges. Propidium iodide (PI) for the visualization of dead bacteria was excited with a 594 nm LED (LS620), 561 nm laser line (Cell Observer) or 552 nm laser line (SP8). The emission of PI was collected within 612-680 nm (LS620), 575-625 nm (Cell Observer and SP8) ranges. The acquisition was performed using 20× dry objectives in a sequence to avoid a spectral bleedthrough. For a given set of experimental conditions (untreated biofilm and biofilms with antiseptics) the same acquisition settings were applied on each system (LED/laser power, exposure time (LS620 and Cell Observer), camera/photomultiplier gain) to enable quantitative comparisons between the conditions. The settings and signal intensity were always set on the brightest samples to avoid oversaturation. Fluorescence intensities were measured in single wide-field images from a focus plane (LS620) or in single optical planes recorded on confocal systems in the Fiji/ImageJ software (NIH, Bethesda, MD, USA). Data are presented as single planes or maximum intensity projections of confocal Z stacks, rendered in the Imaris software (Oxford Instruments, Abingdon, UK). PI is shown in red/orange and SYTO-9 in green color. The denotations “top”, “middle” and “bottom” used to describe the thickness of biofilm structure, are applied in their biological meaning, i.e. (“bottom” – part of biofilm with the direct contact with the plate surface, “middle” – part of biofilm placed over the surface-adhered layer; “top” – the part of biofilm constituting the apexes of hill-like biofilm structure)

### 2.4. Scanning Electron Microscopy Analysis

The biofilms were cultured in the analogical manner as described in Materials and Methods 2.2., with such a difference that the polystyrene coupons were placed in the bottom of the 24-well plate and served as the surface for staphylococcal biofilm growth. Next, the samples were gently cleansed in PBS (Sigma-Aldrich, Darmstad, Germany) buffer as it was described in; fixed in glutaraldehyde [28] (POCH, Wroclaw, Poland) and dried in a critical point dryer EM CPD300 (Leica Microsystems, Wetzlar, Germany). Subsequently, the samples were subjected to sputtering with Au/Pd (60:40) using EM ACE600, Leica sputter (Leica Microsystems,Wetzlar, Germany). The sputtered samples were examined using a scanning electron microscope (SEM, Auriga 60, Zeiss, Germany).

### 2.5. Staining staphylococcal biofilms with Violet Crystal method

The 0.5 McFarland density of the bacteria suspension in TSB medium was prepared and next diluted to 1 × 10^5^ CFU/mL. A total of 1mL of the suspension was added to the well of a 24-well microtiter plate (Wuxi Nest Biotechnology, Wuxi, China) and incubated for 24 h at 37 °C. Subsequently, the non-adhered cells were removed, and the plate was dried for 10 min at 37°C. Next, 1mL of 20% (v/v) water solution of crystal violet (Aqua-med, Lodz, Poland) was added, and the mixture was incubated for 10 min at room temperature. After incubation, the solution was removed, the biofilm was gently washed twice with 100 μL of 0.9% NaCl (Stanlab, Lublin, Poland), and dried for the next 10 min. Next, the image of dyed biofilm was captured photographically.

### 2.6. Impact of rinsing on fluorescence intensity of SYTO-9 and propidium iodine

The procedures were performed as in Material and Methods 2.2. and 2.3. with such a difference that rinsing with water performed once, twice, three or four times. After every rinsing the analysis with use of Etaluma Lumascope 620 fluorescent microscope (San Diego, CA, USA) as performed in Material and Methods 3.3. to capture the changes in Fluorescence Intensity from SYTO-9 and propidium iodine dyes.

### 2.7. Image processing of staphylococcal biofilms using ImageJ software

The captured biofilm pictures were processed using ImageJ (National Institutes of Health, Bethesda, MD, USA) First, RGB pictures were divided into green and red channel sub-images and subsequently changed into 32-bite grey types. Next, the mean grey value was extracted from images of every type. The mean grey value correlates with the value of fluorescence intensity and it is defined as the sum of the values at all pixels divided by the number of pixels. For clarity sake, the OY axes in the Figures dealing with intensity values are presented as Fluorescence Intensity. In case, when application of ABE [%] formula required the assessment of BCA [%] (Biofilm-Covered Area), the images were changed into 32-bite grey type, next the threshold option was applied to differentiate the region of interest (staphylococcal biofilm) from the areas non-covered with biofilm (referred to as the “Background”); subsequently the options Analyze->Set Measurements->Area was applied to calculate percentage of area covered with biofilm.

### 2.8. The assessment of Antiseptic Biofilm Eradication value

The staphylococcal biofilms were cultured as described in Material and Method 2.2., each strain in 6 technical repeats. After medium removal, 1mL of antiseptic (PHMB, NaOCL or PVP-I) or saline (control) was introduced for 1 hour. Next, antiseptics or saline were removed, the universal neutralizing agent (Saline Peptone Water, Biocorp, Warsaw, Poland) was introduced for 5 min. After this time, the neutralizing agent was removed. Then, Live/Dead dye was introduced as presented in Material and Methods 2.3.Twelve fields of view (FoV) in each replicate (in every biofilm-covered well) were imaged. The selection excluded FoVs from the peripheral parts of the well, because of the strong removal of dyed biofilm cells due to shear forces which resulted in very high standard deviations of obtained fluorescence data. Therefore, the FoVs were taken from the center of the well in the following manner: the oval mark was made (using felt tip pen) on the center of the well’s external surface (the most external, bottom part of plate, the one touching incubator) before biofilm culturing and further processing. After sample staining, 4×3 fields of vision were taken (3 fields above the oval mark, 3 fields below the oval mark, 3 fields from the left and 3 fields from the right side of the mark). It provided (4×3) x 6 FoVs. The images were taken with Etaluma Lumascope 620 fluorescent microscope. The estimation of Fluorescence Intensity (measured by Mean Grey Value) and Biofilm-Covered Areas were performed as presented in Material and Methods 2.7. To compare the results of ABE [%] with standard method of antibiofilm activity assessment, the quantitative culturing was performed. The biofilm culturing, antiseptic introduction, neutralization was performed in the exact the same manner as in the case of experiment described for ABE [%] assessment. After neutralization, the 1mL of 0,5% saponin (Sigma Aldrich) was introduced to the wells of 24-well plate and the whole setting was subjected to mechanical shaking for 1 minute to de-attach the biofilm. Subsequently, the serial dilutions of obtained staphylococcal suspension in saline were performed; 100µL of every dilution was cultured on the Columbia Agar plates. The plates were incubated at 37°C for 24h, afterwards the number of staphylococcal colonies was counted.

### 2.9. Statistical Analysis

Calculations were performed using the GraphPad Prism version 7 software (GraphPad Co., San Diego, CA, USA). The normality of distribution was assessed by means of the D’Agostino– Pearson omnibus test. Because all values were non-normally distributed, the Kruskal–Wallis test with post-hoc Dunnett analysis were applied. The results of statistical analyses were considered significant if they produced p-values < 0.05.

## 3. Results

Not only the antiseptic but also the antecedent sample’s preparation (rinsing) and subsequent dyeing procedures are of impact on the biofilm spatial structure. To develop the formula that reliably assess the antiseptic efficacy (and includes impact of rinsing and dyeing), the main characteristics of the staphylococcal biofilm cultured in the *in vitro* setting, was analyzed firstly.

Observed from the aerial perspective, the topography of staphylococcal biofilm in *vitro* resembled a hilly landscape (**Figure 1A**). The spatial concentration of cells within this structure was non-equal (for example lower in parts “1” than in parts “2” and “3”, **Figure 1B**), what was manifested by uneven intensity of fluorescence signal from dyed cells in the particular fields of vision. The staphylococcal biofilm, cultured in the microplate setting, was also of a highly confluent character, i.e. biofilm-forming cells covered basically the whole surface of the plate’s wells (**Figure 1B**). The average level of confluency recorded for the 10 different staphylococcal biofilms, cultured in wells of 24-well plate, was 93.6±5.5% (**Table 1S**).

**Table 1.** The measured thickness of biofilm and the corresponding fluorescence level. A.U. – arbitrary units.

|  |  |  |  |  |  |  |
| --- | --- | --- | --- | --- | --- | --- |
| thickness<br>[μm] | 60.48 | 34.56 | 30.24 | 69.12 | 103.68 | 38.88 |
| fluorescence<br>[SYTO-9]<br>A.U. | 104.7 | 80.5 | 79.2 | 147.6 | 178.6 | 64.9 |

The application of confocal microscopy provided a more detailed insight into the distribution of staphylococcal cells (and their state) within the Z-axis (thickness) of biofilm structure. The staphylococcal biofilms, such as the one presented in **Figure 2**, were of hill-like shape and contained regions of different thickness and of different cells’ density, including areas non-covered with cells (“holes”).

The cellular density of biofilms, along the Z-axis, was higher in their middle (M) parts than in the top (T) and the bottom (B) parts (**Figure 3**). In turn, the cellular density of T and B parts was comparable. Such observation was recorded for 100% (n=10) biofilms formed by analyzed strains (**Table 2S**).

**Tab.2.** Main characteristics of staphylococcal biofilm *in vitro* cultured in this research.

| Parameter(s) | Characteristics |
| --- | --- |
| confluency | very high confluency. Rare areas non-covered with cells (“holes”) |
| shape | the “hill-like” character |
| cellular density | higher in the middle than in the top and bottom part of the biofilms. |
| distribution of “Live” and “Dead” cells | a) high share of “dead” cells in the bottom part of biofilms<br>b) high share of “live” cells in the middle part of biofilms<br>c) top part of biofilm may consist of a relatively equal number of “Live” and “Dead” cells or the share of “Dead” cells significantly exceeds the number of “Live” cells |
| thickness | a) high intra-species and individual variability.<br>b) areas of higher thickness not always display a higher fluorescence level comparing to areas of lower thickness |

**Fig. 3.**
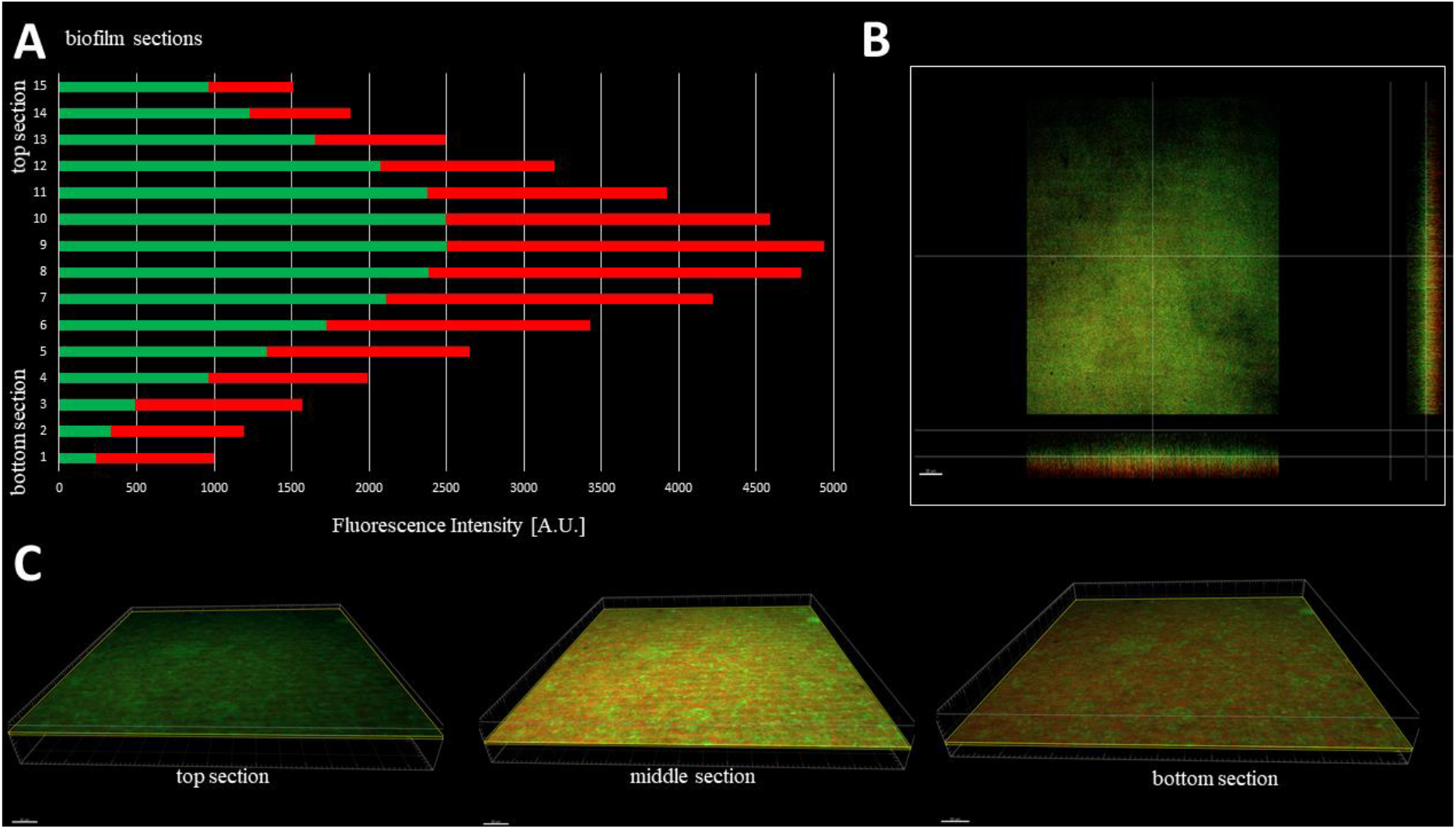
Typical distribution of cellular density through the biofilm. (ATCC 6538 strain) from its top to the bottom section [A, B] and the growing share of dead/cell wall compromised cells in the deeper parts of biofilm [A,C]. Thickness of every section was 2.16µm. Red and green color in [A] represents Fluorescence Intensity of propidium iodide or SYTO-9, respectively; microscope SP8, magn. 40x. Scale bar is 50µm.

With regard to the ratio of altered (damaged) to non-altered (non-damaged) cells within particular parts (B,M,T) of 10 staphylococcal biofilms, the three various patterns were distinguished (**Figure 1S**). In 50% of cases, the majority of altered cells was located in the B part of biofilms, while the majority of non-altered cells in their T part. In case of the second pattern, present in 30% of staphylococcal biofilms, the majority of altered cells was also localized in the B part of biofilm, while the M and T parts consisted of the altered and non-altered cells in various proportions. In case of the third pattern, present in 20% of staphylococcal biofilms, the majority of non-altered cells was localized in the M part of biofilm, while the remaining parts (T, B) of this structure consisted mostly of altered (damaged) cells (**Table 2S**). These types of different cell distribution could also occur simultaneously within a single culture plate well (**Figure 2S**). Next, the areas described in this manuscript as “the holes” (black, non-fluorescent areas pointed as “1s” in **Figure 1B**) were scrutinized. As already showed, the standard *in vitro* culture of staphylococci led to the formation of biofilm of almost full confluency (average above 90%, **Table 1S**). The lowest confluency (81.7%) among the tested strains was displayed by the S1 strain. For the biofilm of this strain, an analysis aiming to evaluate the character of black, non-fluorescent areas was performed using L/D dyeing (**Figure 4**).

**Figure 4.**
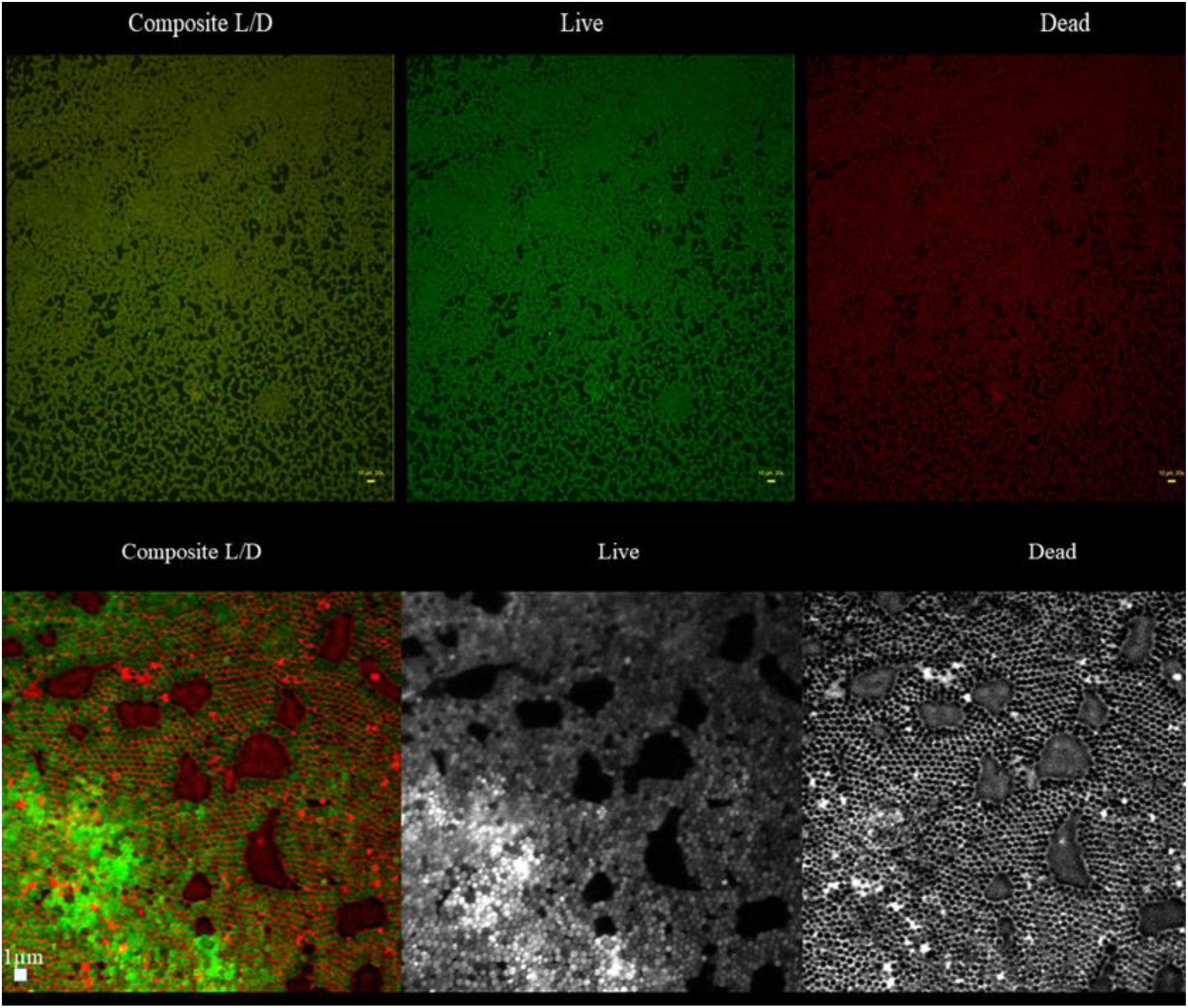
The presence of black non-fluorescent areas (“holes”) in the biofilm of the S1 strain. The upper part of the figure shows holes in the area magnified 20x (the upper panel consists of a composite image (L/D) and split channels (“live” – middle picture; “dead” – right picture). The bar size is 20µm. The “holes” are not-fluorescent(black areas) in any channel in this magnification. In turn in the 100x magnification (lower panel), “holes” take forms of non-fluorescent or red-fluorescent areas. Bar size is 1µm. For higher visibility, in the lower panel, the dye distribution (color fluorescence) was presented in the grey (from white to black) spectrum of colors. Upper panel: microscope LumaScope 600, magn.20x; Lower panel: Microscope Cell Observer, magn.100x.

The analysis revealed that “holes” are areas non-covered with cells or covered with the DNA-containing debris (the distribution of PI within holes was uniform, with no signs of remaining cellular, oval shapes, **Figure 4**). Next, the analyses of biofilm thickness (understood as number of dyed cells along the Z axis) were performed from the point where the fluorescence level started to be detectable (the T part of biofilm) to the point where the fluorescence level was equal to the threshold (value of fluorescence recorded for an un-dyed sample) level, i.e. to the area where biofilm ended and polystyrene surface of the microplate began. The measured thickness of staphylococcal biofilms differed not only between particular strains, but also within a single sample (biofilm cultured in the particular well of a 24-well microplate) (**Figure 5**).

**Figure 5.**
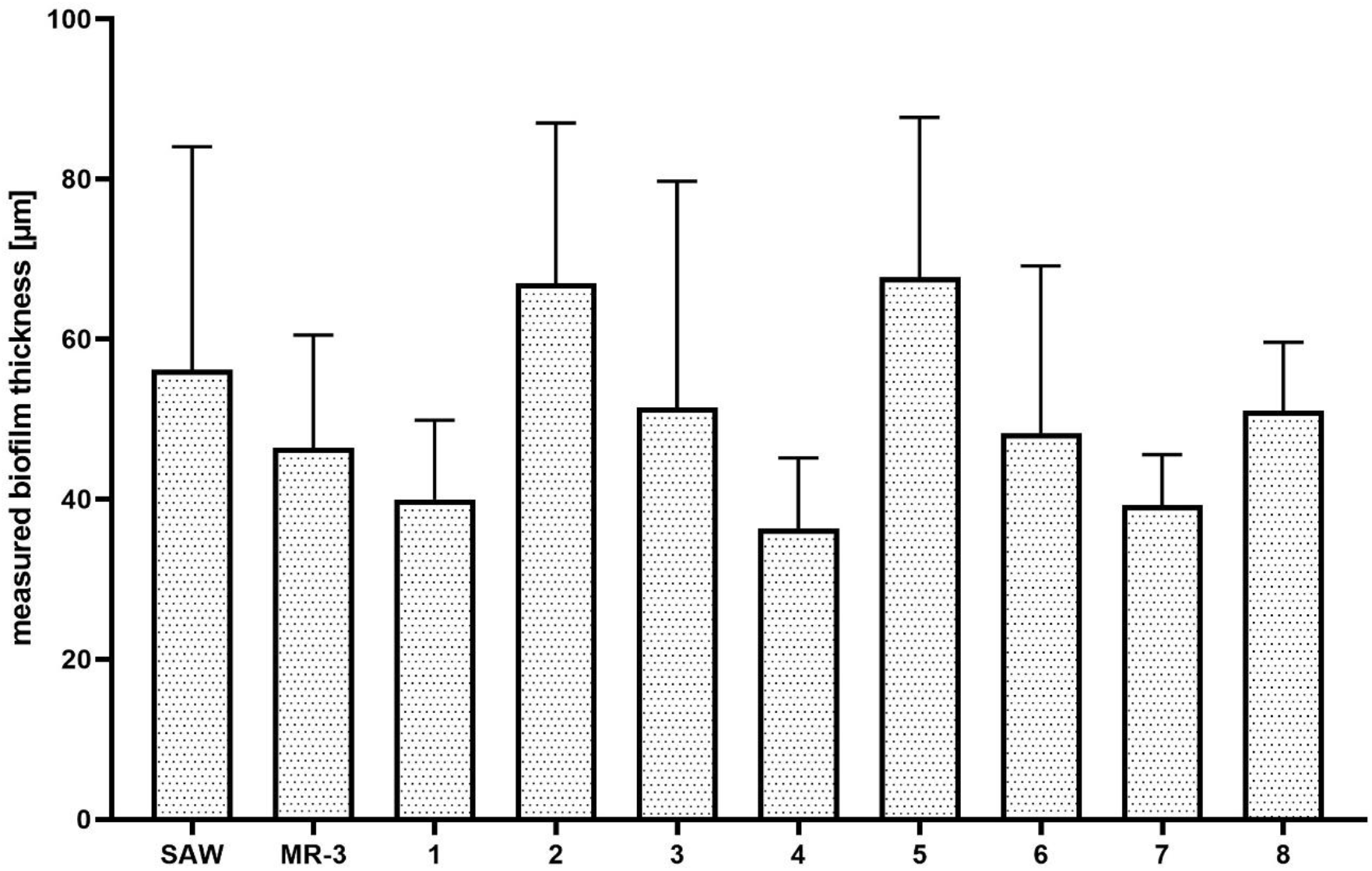
Measured thickness of staphylococcal biofilms (n=10) *in vitro*.

It was also observed that there was no direct correlation between biofilm thickness and intensity of measured fluorescence, which means that specific biofilm could contain the thinner areas where fluorescence level was higher, than is specific, thicker areas (**Table 1**).

The specification of the main characteristic features (thickness, L/D cells ratio, confluency) of staphylococcal biofilms *in vitro*, summarized in Table 2, was necessary to analyze changes of these features after antiseptic introduction and subsequent dyeing process.

The introduction and removal of liquids (rinsing, antiseptic introduction, dyeing) to and from the culture vessels, where biofilms are cultured, belong to the recognized disadvantages of microplate-based methods, as they lead to random de-attachment of biofilm. **Figure 6** presents the impact of a single step (medium removal) on the spatial distribution of the staphylococcal biofilm during 90 seconds of observation. The processing of the initial (15 sec.) and the final image frame (90 sec.) with the use of imageJ software indicated (**Figure 6, lower panel**) that areas covered with cellular structures differed by 23.027%.

**Figure 6.**
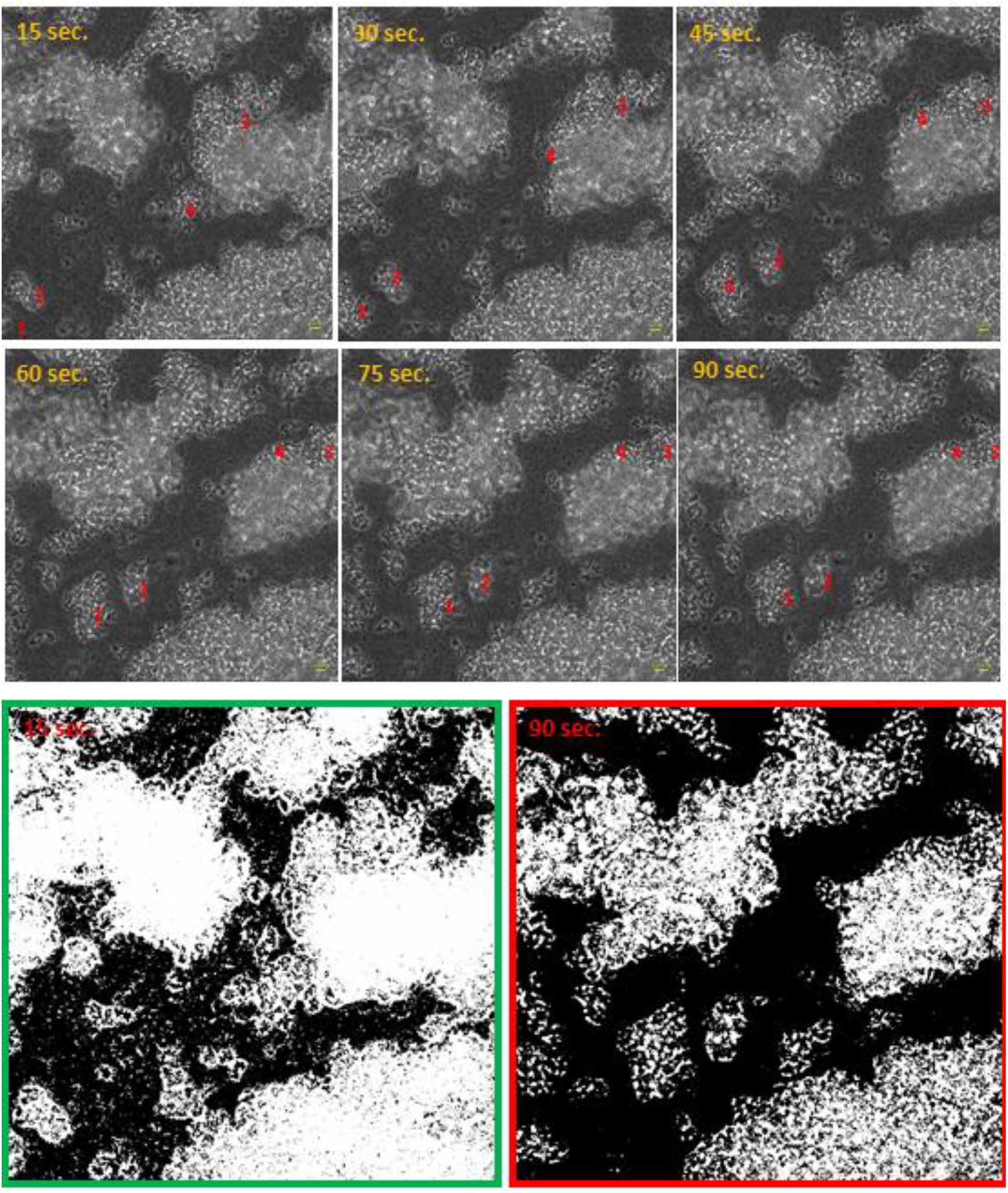
The impact of medium removal on the distribution of staphylococcal (ATCC33591) biofilm within 90 seconds of observation. Numbers 1-4 designate exemplary staphylococcal cell aggregates drifting over time and re-shaping biofilm structure. The lower panel presents changes in staphylococcal biofilm distribution following the implementation of image processing software. The scale bar is 10µm, magn.40x, microscope Lumascope 600. Please click the https://www.youtube.com/watch?v=uwTtHIurDfw&t=24s to watch a video showing the process of biofilm de-attachment.

The process of rinsing is of paramount importance not only due to the impact on the staphylococcal biofilm structure and confluency but also because it correlates with an un-even change in the fluorescence intensity level measured for SYTO-9 (“live”) and PI (“dead”) cells. As can be seen in **Figure 7**, the subsequent rinsing steps lead to a significantly stronger reduction of the PI fluorescence level compared to the reduction in the SYTO-9 fluorescence level.

**Figure 7.**
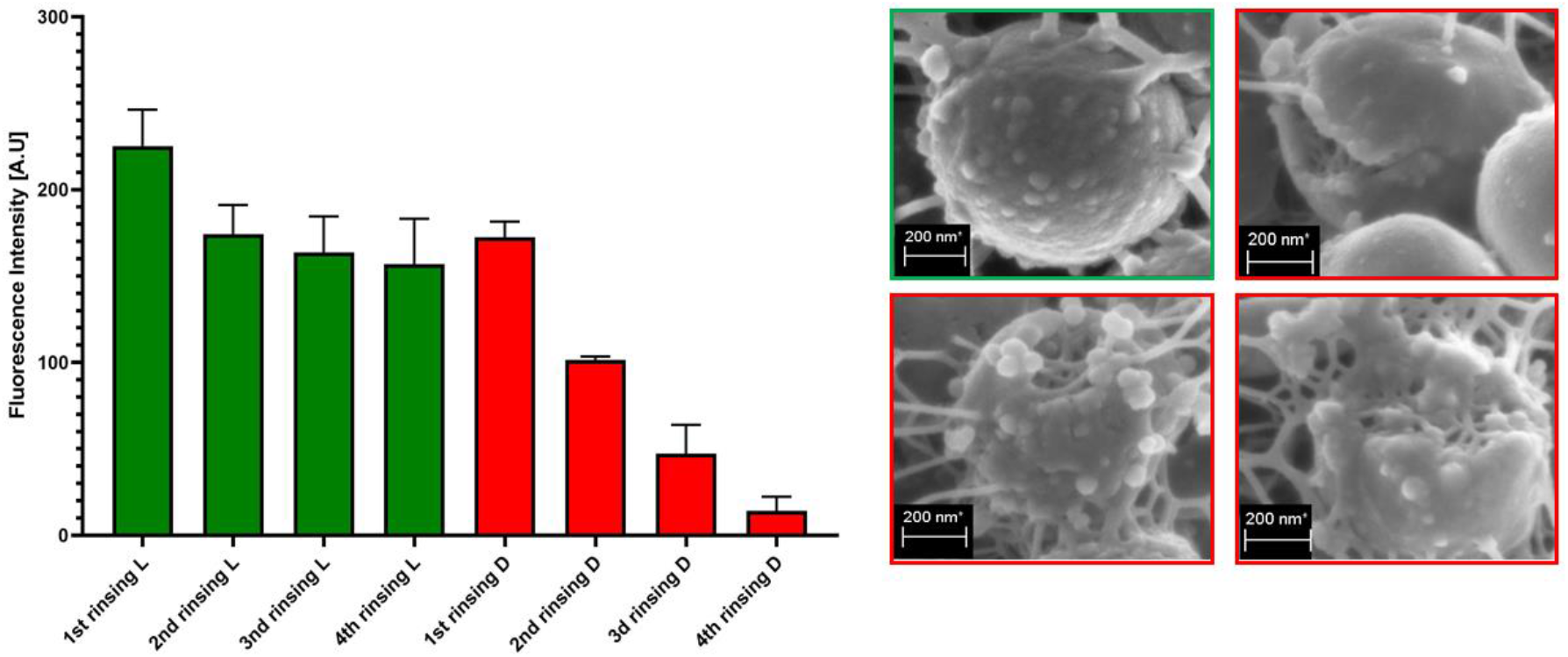
The impact of subsequent rinsing steps on fluorescence intensity of SYTO-9 (L) and PI-dyed (D) biofilm-forming cells of the ATCC6538 strain. The right panel shows staphylococcal cells of an intact cellular wall (green frame) and staphylococcal cells of an increasing level of cell wall damage (red frames) in result of rinsing. The scale bar is 200nm. Magnification 100000x, Microscope Auriga 60.

Another variables, important from the view of coherence of results obtained are of operator-related character as shown in **Figure 3S** and summarized in the **Table 3**. Based on the data collected in **Table 2** and **Table 3**, the proper conclusions can be drawn from the images of non-treated biofilms (control group) and biofilms treated with antiseptic, such as the ones presented in **Figure 8**.

**Table 3.** **Main preparative/process variables associated with the deviation in the obtained results and the suggested, counter-actions**.

| Parameter/process/phenomenon | Characteristics | Action |
| --- | --- | --- |
| rinsing (liquid introduction/removal) | randomly re-shapes biofilm architecture | a) perform with precaution;<br>a) if possible, standardize all sub-steps (place of pipette placement, force of liquid introduction/removal)<br>b) perform preliminary studies to check whether the number of rinsing steps may be reduced without compromising the quality of the performed analyses<br>c) due to rheological phenomena, the rim of the well and approximate region of pipette placement is of low usability for subsequent image processing<br>d) engage a single operator into the entire process of biofilm culturing, antiseptic introduction, and biofilm dyeing |
| PI dye is of lower applicability than SYTO-9 | PI dye is removed from the experimental setting to a higher extent than SYTO-9 | a) consider the application of SYTO-9 only and compare the fluorescence level of SYTO-9-dyed biofilm treated with antiseptic to SYTO-9-dyed biofilm non-treated with antiseptic |

**Figure 8.**
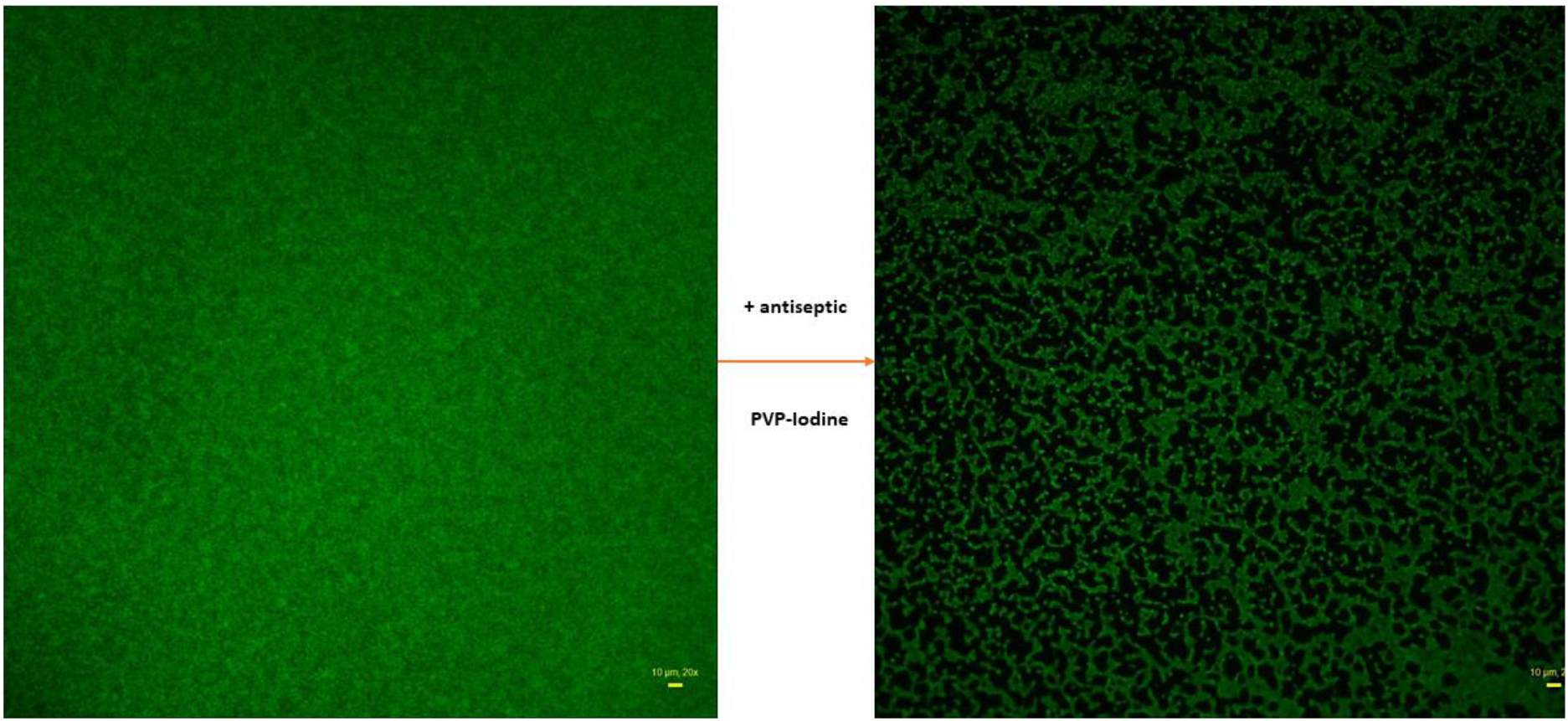
Staphylococcal biofilm of the ATCC 6538 strain non-treated (left picture) and treated (right picture) with an antiseptic (PVP-iodine). Biofilms dyed with SYTO-9 only. Aerial perspective, epifluorescent microscopy. Data collected with wide-field Lumascope 620 magn. 20x, scale bar is 10µm.

The non-treated biofilm (**Figure 8, left picture**) is almost fully confluent, (contrary to the treated-one), nevertheless it contains regions of differentiated cellular density, manifested by differentiated levels of fluorescence within specific regions of observation (i.e. of more and less intense green color).

In turn, the high proportion of image of treated biofilm is covered with black, non-fluorescent areas. The intensity of signal from SYTO-9-dyed cells is lower than in case of non-treated biofilm. Thus, the main differences observed between the two pictures in **Figure 8** are: i) drop in confluency of treated biofilm and ii) drop in the fluorescence level in treated biofilm comparing to the non-treated one. The drop in confluency as well as in the fluorescence level may result from antimicrobial activity of the antiseptic applied, but also, to some extent, from the rinsing procedures. Therefore, the mathematic formula enabling to assess the level of the impact of an antiseptic on staphylococcal biofilm should consist of two components, the first of which should deal with confluency changes, while the second one with the changes in live cells number (measured indirectly with the SYTO-9 dye). Therefore, the first component was described as Biofilm-Covered Area (BCA), assessed as the percentage [%] of culture-well area covered by adherent cells. The exemplary image processing leading to BCA [%] calculation by means of ImageJ is presented in **Figure 9**. As an effect of this action, only areas above an established threshold value are considered to contain biofilm-forming cells, while the remaining areas are considered cell-free (the “holes”, non-fluorescent areas).

**Figure 9.**
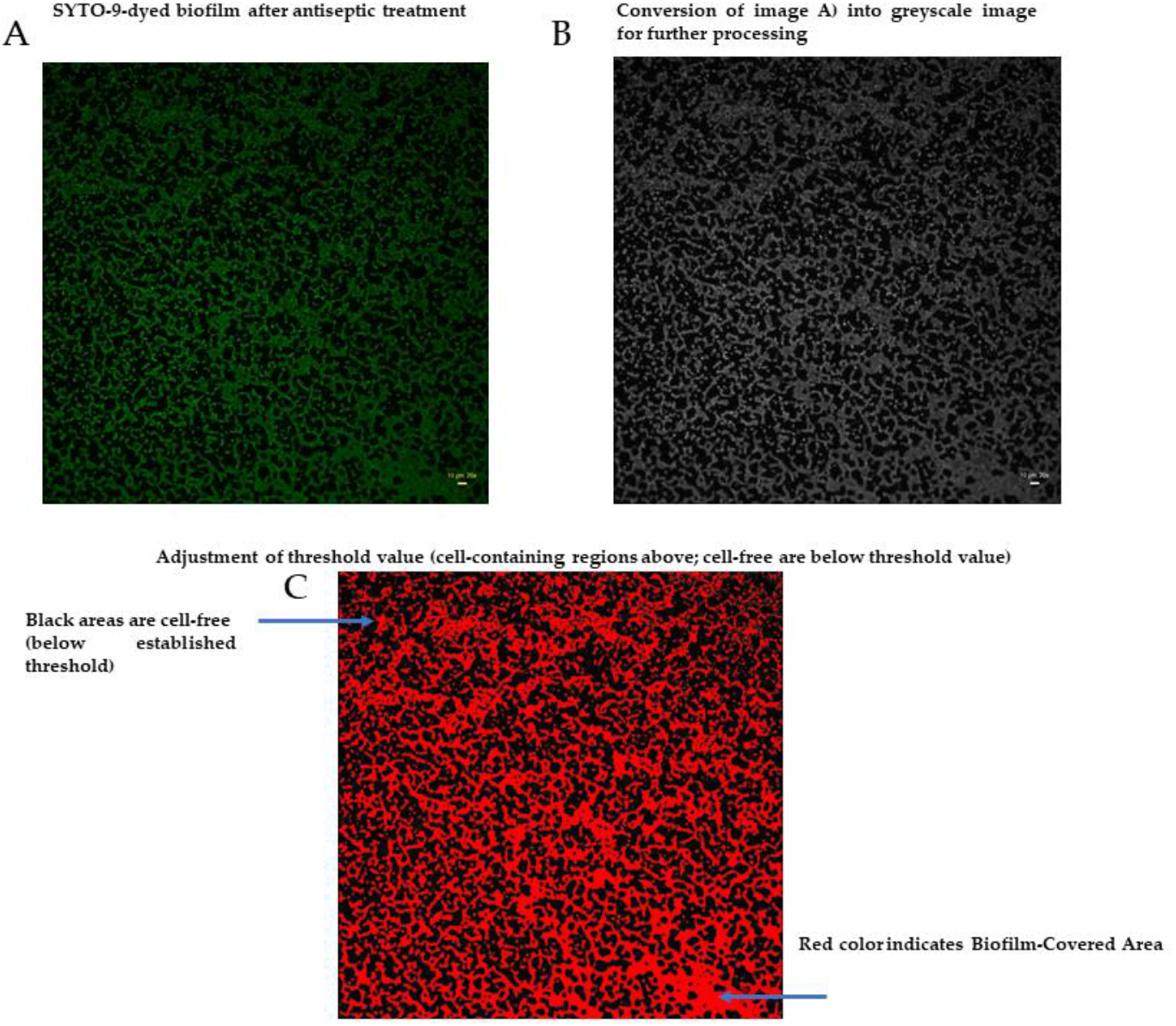
Processing of biofilm image to extract the value of Biofilm-Covered Area [%]. A – biofilm after treatment with antiseptic; B,C – processing stages aiming to extract the value of BCA [%].Data collected with wide-field Lumascope 620, 20x, scale bare is 10 µm.

The BCA [%] calculated for the biofilm treated with antiseptic (**Figure 8, right picture**) was 49%, and 99% for the biofilm treated with saline, (**Figure 8, left picture**). Thus, the antiseptic activity correlated with a drop in BCA value by 50% (99%–49%). The second parameter introduced to the formula was a drop in cell number in the BCA, referred to as the Biofilm Fluorescence Intensity Drop (BFID). It indicates the antiseptic-caused death of a certain number of cells (but not all of them) within specific Z-axis (thickness) of the biofilm. To calculate the BFID, the FI of treated biofilm-covered areas was compared to the FI of non-treated biofilm areas. The BFID is calculated only for Biofilm-Covered Area, not for black, non-fluorescent regions. Following the example of biofilms presented in **Figure 8**, FI of BCA of treated biofilm and non-treated biofilm (BCA regions) was 44.99 vs. 104.38, respectively. It means that FI in antiseptic-treated BCA constituted a fraction of 0.43, i.e. 44.98/104.38 of the non-treated biofilm. The BCA value of treated biofilm was 49%, so the fluorescence level in this area was decreased by 21.07% (0.43×49%). Therefore, the antiseptic activity against this not completely eradicated biofilm can be expressed as the BFID equal 49%–21.07% = 27.93%.

Combining the BCA and BFID components, the following formula of Antiseptic’s Biofilm Eradication (ABE [%]) was achieved:

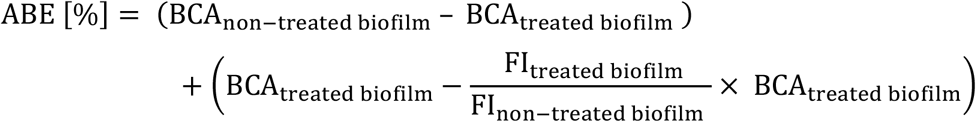

and after simplification:

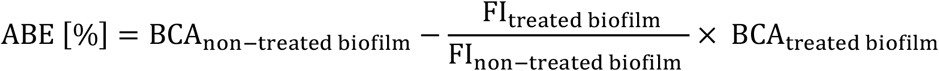

Therefore, the ABE [%] for the exemplary staphylococcal biofilm treated with PVP-I (**Figure 8**) was 78.77%.

Having the ABE [%] formula established, the assessment of the effect of exemplary antiseptics (PHMB, PVP-I and NaOCl) on staphylococcal biofilm of all 10 scrutinized strains was performed (**Figure 10**).

**Figure 10.**
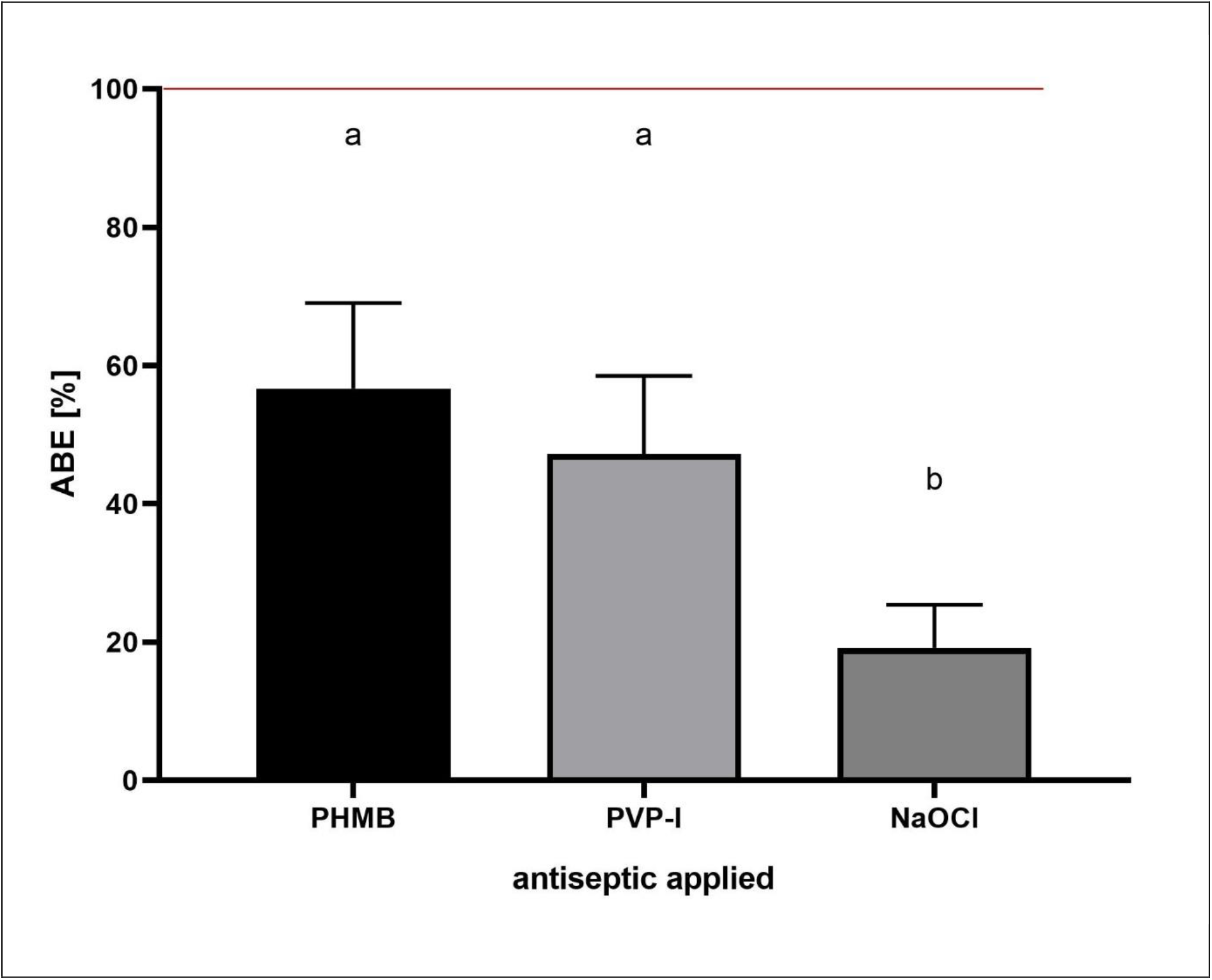
ABE [%] value. recorded for polyhexamethylene (PHMB), povidone-iodine (PVP-I) and hypochlorite solution (NaOCl) towards staphylococcal biofilms (n=10) formed *in vitro*. ABE [%] – Antiseptic’s Biofilm Eradication. Letters a,b show statistically significant differences in ABE [%] values between PHMB and PVP-I vs. NaOCl. The red line represents the level of formation of non-treated, control biofilm in relation to which the loss of biofilm-covered area and fluorescence intensity in treated biofilms are calculated.

The results presented in **Figure 10** show comparable antibiofilm activity of PHMB and PVP-I, and a statistically higher (p>0.5) activity of these compounds in comparison to the NaOCl activity. Finally, the same experimental setting as the one presented in **Figure 10** was performed, but the antiseptics’ antibiofilm activity was assessed using standard, quantitative culturing (QC) method. The [%] biofilm eradication was 48±13%, 44±5% and 10±3% for PHMB, PVP-I and NaOCl, respectively. The results obtained by both methods (ABE% and QC) were cohesive, showing comparable level of eradication after use of PHMB and PVP-I and low level of eradication after use of NaOCL. The differences in eradication level recorded after use of ABE% and QC methods were 8,3%, 2,7% and 9% for PHMB, PVP-I and NaOCL, respectively.

## 4. Discussion

The biofilms can be perceived as highly differentiated biological systems, able to efficiently adapt to the specific conditions in human organism **[14]**. Therefore, to assess the impact of antimicrobial agents on biofilm, the differentiated techniques, measuring changes in biofilm’s crucial components, should be applied. With this regard, the culturing of biofilm-forming cells and metabolic activity assays provide data of respectively quantitative and semiquantitative character, while microscopic images of biofilms tagged with fluorescent dyes are presented frequently to back-up the aforementioned results **[15]**. The aim of this line of investigation was to use the possibilities offered by the software for image processing and to provide quantitative data on antibiofilm impact of antiseptic agents using microscopic images of biofilm dyed with reagents differentiating live cells from the dead ones (“Live/Dead”). To obtain this aim, the features of staphylococcal biofilm, cultured *in vitro*, were firstly analyzed. The data presented in the **Figures 1-5** can be summarized as follows: the staphylococcal biofilm *in vitro* is a cellular structure of differentiated thickness and of highly confluent character. The rare gaps (“holes”) in the continuity of the biofilms are devoid of cells and/or they contain cellular debris. The bottom and the top parts of biofilm display lower cellular density than the medium parts of biofilm. The majority of dead cells in staphylococcal biofilm *in vitro* are localized in the bottom part of this structure, nevertheless some of the biofilms displayed also the pattern in which dead cells prevailed not only in the bottom, but also in the top parts. Such staphylococcal biofilm heterogeneity may be of high importance with regard to the activity displayed by these antiseptics which act through disintegration of microbial cell wall (for example polyhexamethylene biguanidine **[6]**). One may expect that their effectiveness may differ depending on the type of biofilm they will be used against, i.e. whether the antiseptic molecules will have an initial contact with the heavily damaged cells of such biofilm as presented in **Figure 1S, pattern C** or with mostly non-compromised cells forming biofilm structure presented in **Figure 1S, pattern A**. In fact, the normative methods of assessment of antiseptic activity **[16]** imply the use of an “organic burden” (bovine serum albumin and/or blood cells) to analyze the impact of this load on antiseptic molecules activity, which is mostly negative (the effectiveness drops). One may hypothesize that the T part of biofilm presented in **Figure 1S, pattern C** may act in a similar manner as organic load applied in normative methods. The antimicrobial activity of most antiseptics relies on destruction of bacterial cell walls (and biofilm matrix) and one may hypothesize that the result of such activity may be empty spaces in the biofilm structure, contributing to the drop in confluency level. Such a drop in confluency is commonly observed in eukaryotic cell cultures *in vitro*, after the introduction of molecules displaying high cytotoxic potential **[17]**. The occurrence of holes (black, non-fluorescent areas) in biofilm, would also suggest that the specific antiseptic was able to penetrate through the whole biofilm structure (from part T to B along the Z-axis and to kill biofilm-forming cells). The high tolerance of the deeper parts of biofilms to antibiotics and the immune system components is one of the main clinically-recognized challenges impeding effective biofilm’s eradication **[18]**. Therefore, these antiseptics whose activity correlates with the formation of new holes within biofilms should be considered the most effective ones. The analysis of the properties of staphylococcal biofilm *in vitro* should also include another factor, namely the height (thickness) of the biofilm, understood as the number of subsequent cells along the Z-axis. The parameter of thickness is crucial with regard to the application of antiseptics against biofilm, because it may be hypothesized that the application of cell wall-destructing antiseptics should lead to the removal of cells and to a decrease in biofilm thickness. This assumption is valid not only because of the aforementioned activity mechanism of these antiseptics, but also because of the rheological phenomena occurring during antiseptics’ introduction, removal and subsequent laboratory stages of dye introduction and microplate well rinsing. As was observed, the measured thickness of staphylococcal biofilms differed not only between particular strains, showing explicitly the importance of intra-species variability **[19]**, but also within a single sample (well of a 24-well microplate). This thickness differentiation is meaningful, taking into consideration the activity of antiseptics against biofilm. Firstly, the process of antiseptic’s pouring in initiates a number of rheological and dynamic processes resulting in (random from our perspective) breaking off of biofilm parts. Secondly, the level of biofilm eradication depends on the specific antiseptic agent’s mode of action, its ability to penetrate through the matrix and on the cells’ susceptibility. It may be hypothesized that the thinner the biofilm in a particular area, the higher probability that the activity of an antiseptic will result in higher biofilm removal and formation of holes. Nevertheless, as shown in **Table 1**, the thickness of biofilm does not always correspond to the cellular density of cells measured by fluorescence level. The attempt of explanation of this phenomenon is presented in **Figure 11**.

**Figure 11.**
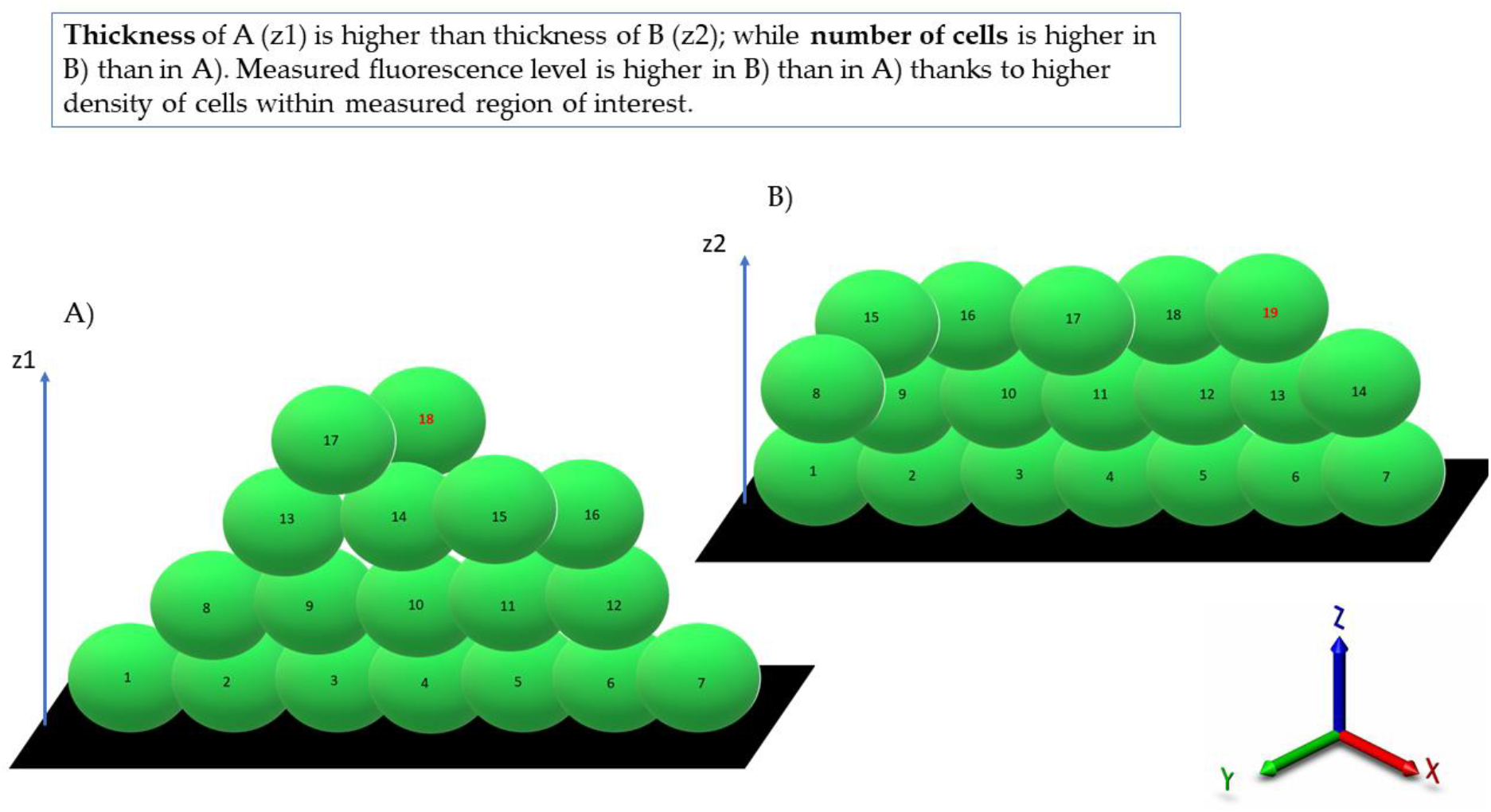
An attempt at explanation of the data presented in Table 1. Because of various cellular density between A) and B) parts of biofilm, the level of fluorescence of biofilm B) is higher than that of biofilm A), although biofilm A) is thicker (4 “layers”) than biofilm B), which consists of 3 “layers”. z1, z2 – Z-axis showing biofilm thickness; green oval shapes – SYTO-9-dyed staphylococci emitting fluorescence; numbers in the central points of oval shapes show a higher number of cells in biofilm A) in comparison to biofilm B).

Following this thought, these thinner parts of biofilm, which display high cellular density, may be harder to eradicate than biofilm that is thicker but consists of a lower number of cells. The above-mentioned considerations are valid if we assume that the level of emitted fluorescence is the same (or at least approximately the same) for every cell in the biofilm, regardless of its physiological state and spatial location within the biofilm. Unfortunately, the performance of an experiment aiming to check the aforementioned possible differences is far beyond the technological possibilities offered by contemporary technology.

After analysis of crucial features of the staphylococcal biofilm, in the next step of investigation, the determination of the impact of liquid introduction/removal to the experimental setting was performed. With regard to the images and quantitative data obtained by means of confocal/epifluorescent microscopy, the de-attachment of biofilm during procedure of rinsing is of paramount (and negative) importance, because it leads to the formation of random holes in the biofilm structure which may be interpreted as resulting from the eradicative force of the introduced antiseptics. The standard procedure of analysis of antiseptic impact on biofilm *in vitro* includes at least 11 steps of liquid introduction and removal. These are: i) aspiration of medium, ii) introduction/removal of rinsing liquid, iii) introduction/removal of antiseptic agent, iv) introduction/removal of antiseptic neutralizer, v) introduction/removal of L/D mixture, vi) introduction/removal of rinsing liquid). The number of steps may be higher, because specific protocols recommend to repeat specific steps (mostly ii and vi) twice or three times. Based on the data shown in **Figure 3S** it can be stated that the alteration of biofilm structure when all these steps are applied (bearing also in mind the fact that one of them, antiseptic agent introduction, leads to the destruction of cells) are of a massive and rather unpredictable, character. In our opinion it is one of the most important variables during the whole process of assessment of antiseptic agent impact on staphylococcal biofilm *in vitro*. The fact that some research teams recommend rinsing the plates with tap water or shaking the plate out over a waste tray **[20]**, additionally and rather strongly impedes the obtaining of repeatable and conclusive results. Nevertheless, the processes of liquid introduction and removal cannot be omitted, if L/D dyeing is to be performed. The solutions aiming to decrease, to some extent, the negative impact of rinsing on the reproducibility of results involve: i) gentle aspiration and removal of liquid during manual pipetting, ii) placing the tip of the pipette in the same position (preferably on the rim of the plate’s well), iii) if possible, reducing the number of rinsing, and, last but not least, the performance of an appropriate number of technical repeats and independent experiments (performed in accordance with the general methodology of biological quantitative experiments). The process of rinsing is of paramount importance not only due to the impact on the staphylococcal biofilm structure and confluency but also because it correlates with an un-even change in the fluorescence intensity level measured for SYTO-9 (“live”) and PI (“dead”) cells. This statement is one of the main discoveries made in the present study because it re-defines the usability of the last-mentioned dye for the analyses of the impact of antiseptic agents on staphylococcal biofilm. As can be seen in **Figure 7**, the subsequent rinsing steps lead to a significantly stronger reduction of the PI fluorescence level compared to the reduction in the SYTO-9 fluorescence level. This observed fluorescence intensity drop could be also due to the phenomenon referred to as the photobleaching as well as due to the lower photostability of PI comparing to SYTO-9 **[21]**. It may be also hypothesized that PI binds to damaged/compromised cells (examples of such cells are presented in the right panel of **Figure 7**), which are more vulnerable to be flushed out by the introduced liquid. The general concept of application of these two dyes is based on the idea that the activity of an antimicrobial should correlate with a drop in fluorescence level from SYTO-9 dyed cells with a simultaneous increase in the fluorescence level from PI-dyed cells. However, the observation from this study (**Figure 7**) explicitly shows that PI is of lower usability for quantitative analyses of staphylococcal biofilm *in vitro* than the SYTO-9 dye. In our opinion, the application of PI may be valid for images of a high aesthetic value, while the quantitative measurement of the viability of staphylococcal, biofilm-forming cells should be performed with the use of SYTO-9 only (by comparison of the levels of SYTO-9 fluorescence between biofilm treated with antimicrobial and untreated, control biofilm). The above statement is particularly valid with regard to these antiseptic agents which disintegrate the continuity of staphylococcal cell walls. In turn, the verification of the discovery presented in **Figure 7** with regard to bacteriostatic agents undoubtedly requires an experimental confirmation. Noteworthy, the proposal of the application of only SYTO-9 for measuring the drop in biofilm’s viability resembles the methodology applied in normative analyses of eukaryotic cell lines (cytotoxicity tests, for example), where the drop in live cells (dyed with tetrazolium salts) is measured after biocide introduction and compared to the un-treated control **[22]**. Moreover, within biofilm structure, a certain population of cells is considered to be “viable, but non-culturable” (VNBC; such cells display a higher resistance to antimicrobials than viable, culturable cells). Because SYTO-9 binds to the VNBC cells, the measured level of fluorescence in the untreated control may include also fluorescence from these cells. On the other hand, such cells are still considered “infective” (and able to cause for example, aseptic loosening of an implant **[23]**), so their detection as part of infective biofilm should be considered valid. A schematic explanation of the phenomena occurring during the introduction of antiseptics, summarizing data collected in this research and resulting in specific types of images from confocal or epifluorescent microscope, is presented in the upper and lower panel of **Figure 12**, respectively.

**Figure 12.**
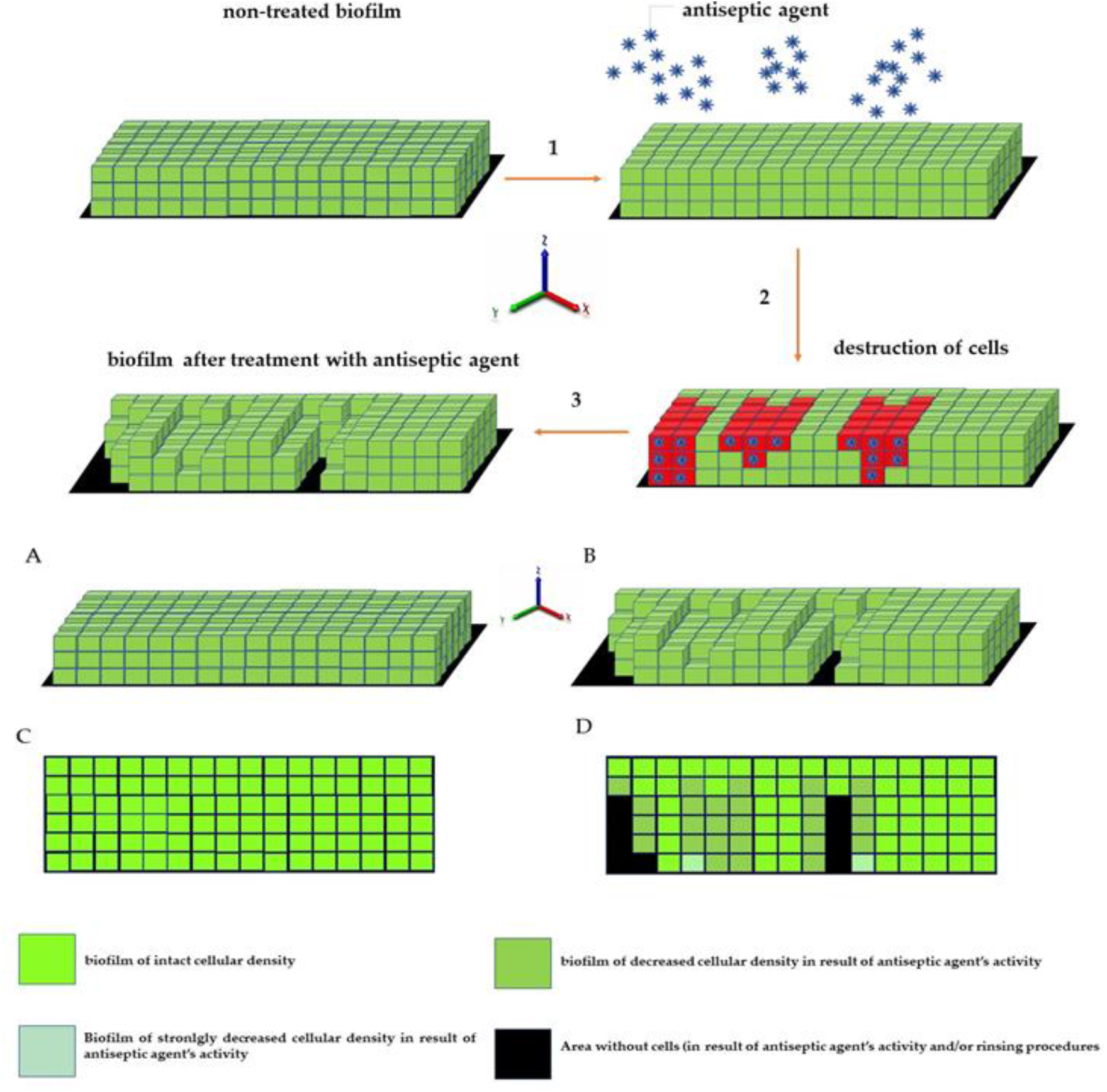
Schematic presentation of the phenomena occurring during antiseptic activity against biofilm reflected in the formation of “holes” and in a drop in fluorescence intensity. For picture clarity sake, the staphylococcal cells are shown as cubes, while intact biofilm formed by the cells is presented as an even (with regard to height) structure. 1) – introduction of antiseptic agent to biofilm; 2) antiseptic agent activity; 3) destruction of cells (resulting from antiseptic agent activity) and removal of cells (resulting from antiseptic agent activity and rinsing). A,C -intact biofilm in 3D and -xy plane, respectively; B,D – biofilm after the application of antiseptic in 3D and -xy plane, respectively.

Based on the data obtained in the course of this investigation line, the mathematic formula, referred to as the ABE [%] was elucidated. The **Figure 10** shows its implementation to assess antibiofilm activity of three commonly applied antiseptic agents – PHMB, PVP-I and NaOCl. By means of ABE [%], comparable antibiofilm activity of PHMB and PVP-I, and a statistically higher (p>0.5) activity of these compounds in comparison to the NaOCl activity was shown. Such results confirm the methodological usability of ABE [%], as they stay in line with the data previously presented by our and other teams concerning the activity of these 3 wound antiseptics **[24,25,26]**; including the lack of effectiveness of NaOCl, containing low, 80 ppm hypochlorite content, showed already by Severing et al. **[13]**. Thus, by research performed in this study, not only a new tool to assess the antiseptic activity against staphylococcal biofilm (ABE[%]) was developed, but also, on a general level, another data-set confirming the usability of microscopic analysis combined with image processing in studies on antiseptics was provided. Although our research may be considered an important step towards complex analyses of antiseptics’ activity against staphylococcal biofilms, we are also aware of certain disadvantages of our work. Firstly, to keep the number of variables under control, basically only 24h-old biofilms were analyzed. If “younger” and “older” biofilms were included to the investigation line, also other patterns of cellular density, biofilm thickness and L/D-dyed cell spatial composition would be probably detected. Secondly, just one type of culturing medium (Tryptic Soy Broth) was applied, while high differences in biofilm structures formed by the same strain, but cultured in different media, were already observed **[27]**. Thirdly, only polystyrene surface was used to culture biofilms, while the impact of various surface types on biofilm structure is undisputable and proven **[28]**. As mentioned, our aim was to keep the number of variables under control, therefore analyses were performed in one particular setting (24h biofilm, TSB medium, polystyrene surface). The implementation of additional variables would translate into overload with data. Still, we are convinced that the range of already performed analyses indicates the directions of subsequent paths that may be explored by other research teams with an aim to increase our knowledge of the phenomena occurring when staphylococcal biofilm is exposed to an activity of locally acting antiseptic agents. Taking into account the fact that *in vitro* results of the efficacy of antiseptic agents against staphylococcal biofilm are frequently applied to back up their use in hospitals and ambulatory units, our work should be considered an important tool providing reliable, quantitative data with this regard.

## Conclusions

**The staphylococcal biofilm *in vitro* displays confluent structure of differentiated thickness and ratio of live to dead cells. These differences depend on intraspecies features**.

**The preparation procedures (rinsing, especially) for microscopic analysis and further image processing significantly alter biofilm structure and have impact on analyses related with application of antiseptic agents**.

**The SYTO-9 dye represents higher value for quantitative assessment of antiseptic impact on biofilm than propidium iodide (PI)**.

**The developed and scrutinized in this research formula of Antiseptic’s Biofilm Eradication, may be of high applicability in assessment of antiseptic activity against 3-dimensional biofilm structure**.

## Funding

This research was funded by National Science Center (Grant No. 2017/27/B/NZ6/02103)

## Institutional Review Board Statement

not applicable

## Informed Consent Statement

not applicable

## Data Availability Statement

the data-set and raw images applied to perform the image panels presented in this study are available in public repository FigShare under the address provided: https://figshare.com/articles/dataset/Data_and_Image_Set/20391495

under the license CCBY 4.0.

## Acknowledgments

AJ would like to thank Howard Bloom (N.Y.C.) for teaching him to always ask: “What do I actually see here?”

## Conflicts of Interest

The authors declare no conflict of interest

## Notes

### Competing Interest Statement

The authors have declared no competing interest.

### Summary of Updates

x

