## Supplementary material for "The efficacy of antiseptic agents against staphylococcal biofilm can be quantitatively assessed with the use of processed microscopic images": ST1

**Table S1. Confluence [%] of analyzed staphylococcal biofilms (number of strains = 10)**

| strain description | confluence [%] | average confluence [%] for n=10 strains |
|---|---|---|
| ATCC 6538 | 99,9 | 93,61±5,55 |
| ATCC 33591 | 99,9 | |
| S1 | 81,7 | |
| S2 | 95,7 | |
| S3 | 94,4 | |
| S4 | 97,8 | |
| S5 | 95,3 | |
| S6 | 91,2 | |
| S7 | 90,8 | |
| S8 | 89,4 | |

**Table 2S. The Fluorescence Intensity of live and dead (L, D, respectively) staphylococcal biofilms with regard to their location** in **T** (top), **M** (middle) and **B** (bottom) parts of biofilm

| **Strain no** | **Part of biofilm** | **Fluorescence Intensity L:D** | **Matches pattern FI M > FI T and FI B?** |
|---|---|---|---|
| **ATCC 6538** | T | 980: 710 | **Yes** |
| | M | 2499:1441 | |
| | B | 258: 742 | |
| **ATCC 33591** | T | 492: 938 | **Yes** |
| | M | 891: 689 | |
| | B | 161:591 | |
| **S1** | T | 664:523 | **Yes** |
| | M | 1725:1705 | |
| | B | 326:623 | |
| **S2** | T | 1353:779 | **Yes** |
| | M | 4130:3174 | |
| | B | 703:1021 | |
| **S3** | T | 954:627 | **Yes** |
| | M | 4265:3094 | |
| | B | 498:295 | |
| **S4** | T | 769:412 | **Yes** |
| | M | 1900:621 | |
| | B | 219:492 | |
| **S5** | T | 205:474 | **Yes** |
| | M | 432:381 | |
| | B | 176:321 | |
| **S6** | T | 522:634 | **Yes** |
| | M | 1421:839 | |
| | B | 283:425 | |
| **S7** | T | 405:174 | **Yes** |
| | M | 571:331 | |
| | B | 108:273 | |
| **S8** | T | 411:290 | **Yes** |
| | M | 710:629 | |
| | B | 138:305 | |

**Fig 1S. Three types (A,B,C) of distribution of dead/cell wall compromised cells vs. viable/cell wall non-compromised cells in staphylococcal biofilm *in vitro***; the vertical cross-section through the biofilm structure. A – strain ATCC 6538; B – strain S1, C -ATCC 33591. Scale bar is 30 µm. Microscope SP8, magn.40x.

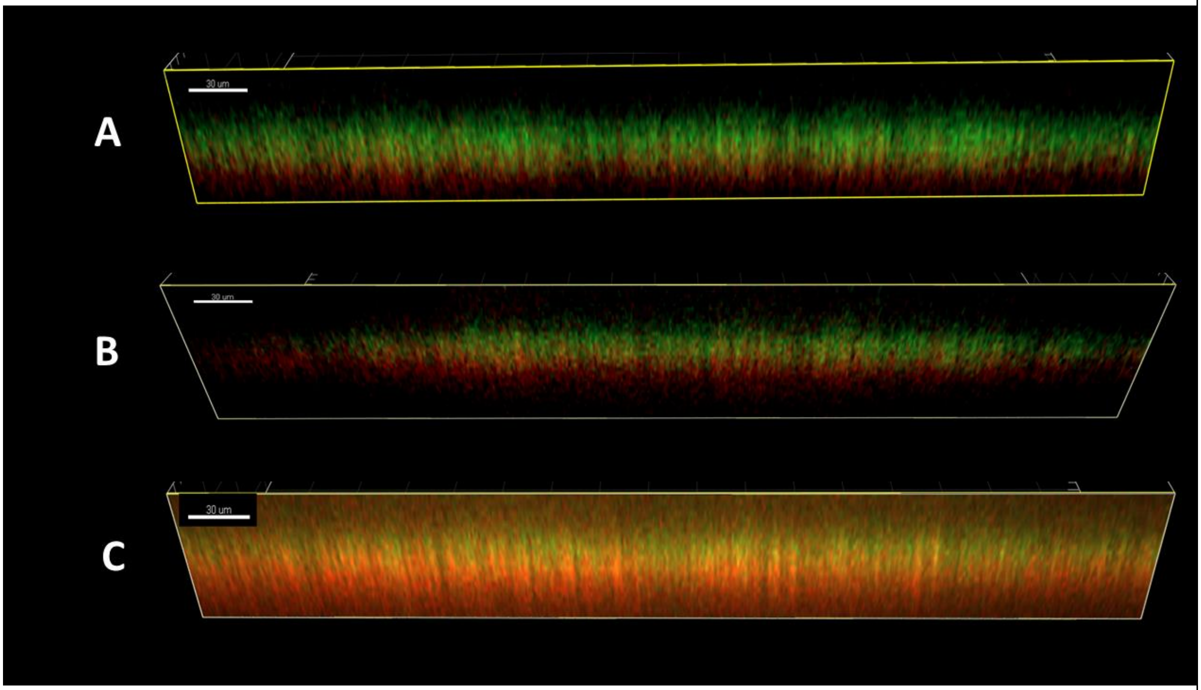

**Figure 2S. The different types of Live/Dead cells' distribution in biofilm within single plate**.
Strain ATCC 33591. Scale bar is 40µm

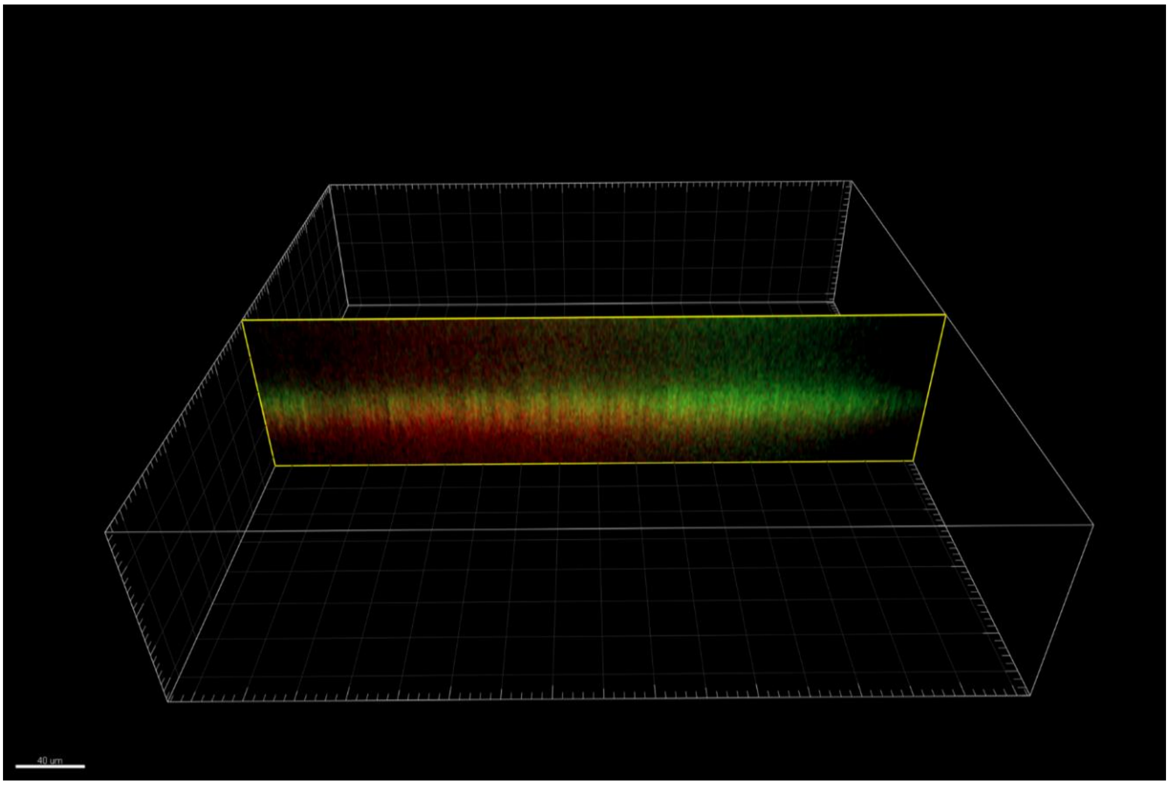


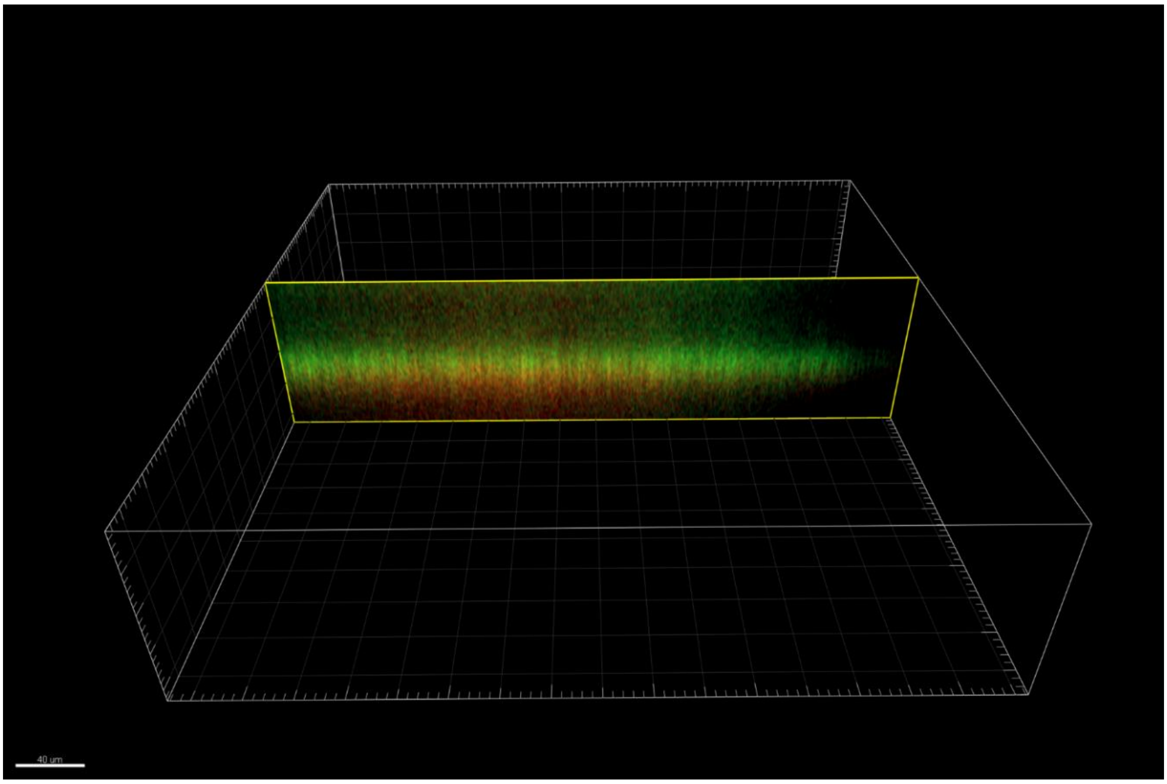

**Figure 3S. The impact of operators' various pipetting habits (A,B,C) on dye distribution in the well (of a 24-well plate) covered with biofilm formed by the same staphylococcal strain (ATCC 6538).** Picture D presents the biofilm of the aforementioned strain dyed with the L/D method by an operator with 3 years of experience in staphylococcal biofilm culturing and dyeing. The red cross indicates the approximate place of pipette tip placement.

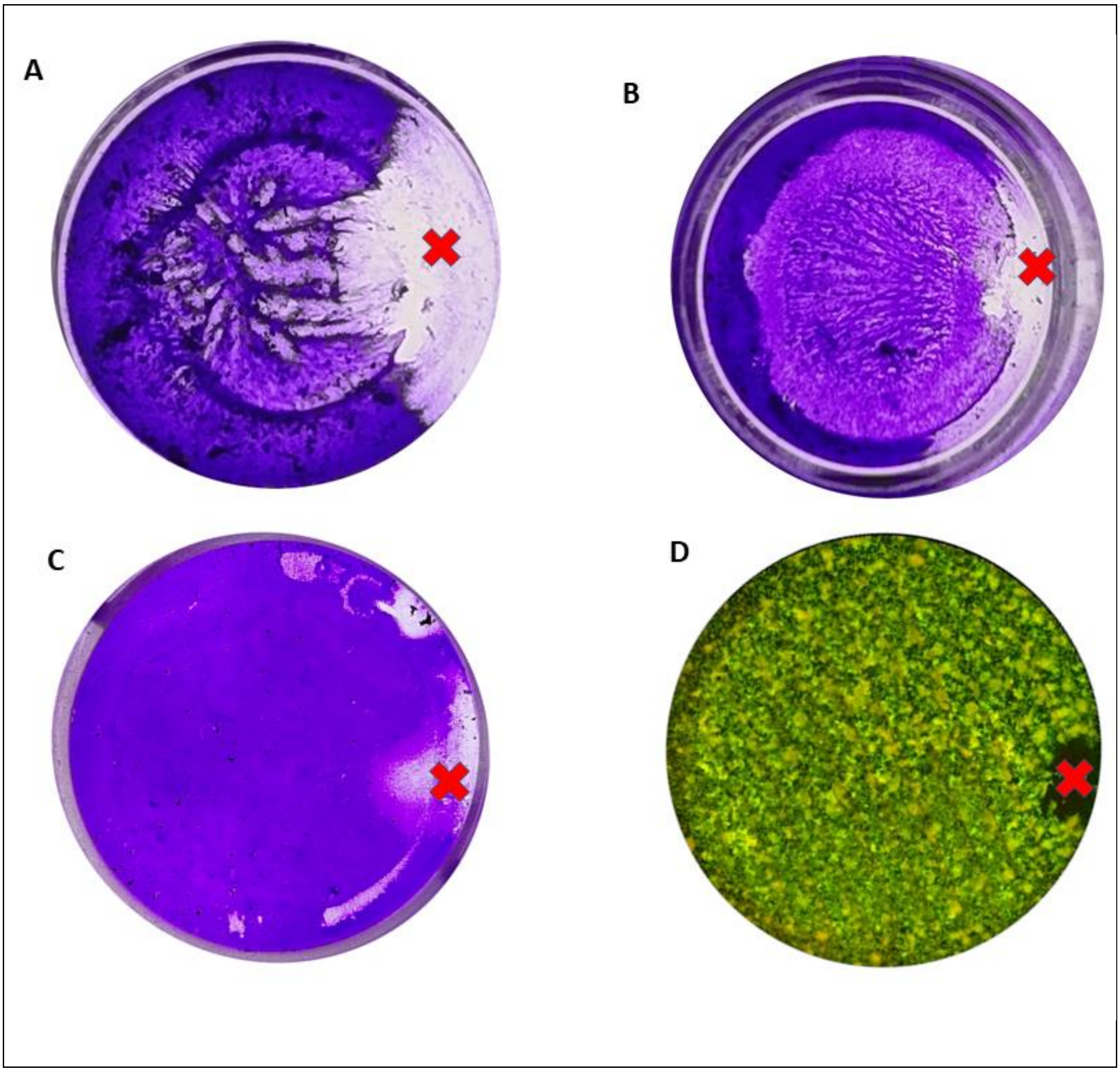
